## Supplemental Methods and Results for "Connectivity in Large-Scale Resting State Brain Networks is Related to Motor Learning: a High-Density EEG Study"

### Supplementary Methods

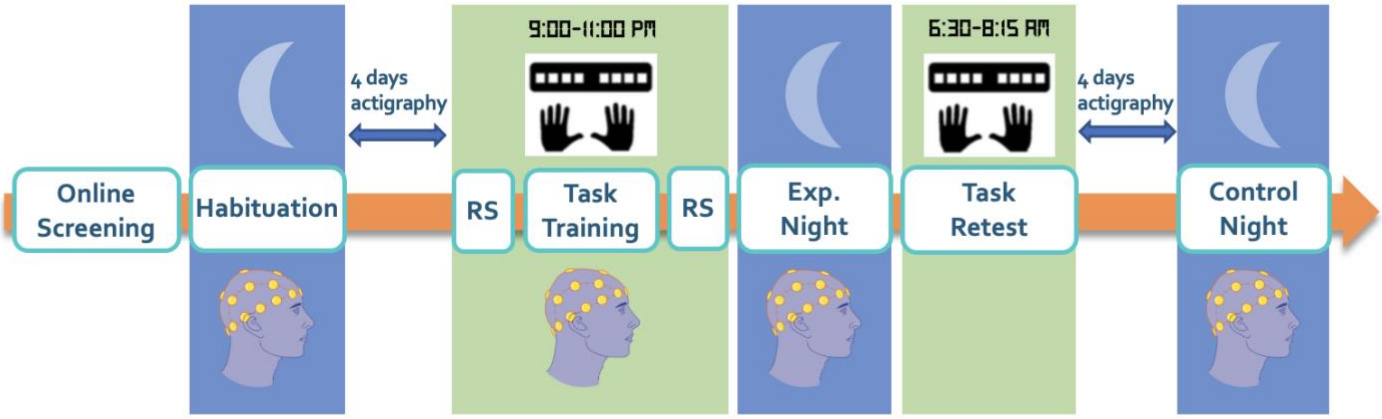


Supplementary Figure 1. Full Study Design. Participants completed three overnight sleep EEG recordings in the lab, and on a separate day also underwent a T1 anatomical MRI scan and a resting state fMRI scan. The first overnight recording was used to habituate participants to sleeping in the lab environment and wearing the hdEEG equipment. The subsequent two nights were either the control night – in which participants slept normally without any intervention – or the experimental night, where they completed the motor sequence learning (MSL) task before sleeping (training) and 45 minutes after waking the following morning (retest). The order of the sessions (i.e., experimental vs. control) was counterbalanced across participants.

**Serial Reaction Time Task**:

A random SRT task [1] was implemented using Matlab Psychophysics Toolbox v. 3, in order to assess general motor execution. During the task, eight squares were visible on the screen, with each square corresponding to one of the keys on a specialized keyboard, and to one of the eight fingers used (excluding thumbs). The outline of the squares alternated between red (rest) and green (practice). During the practice blocks, a green square appeared on the screen in one of the 8 locations outlined, and the participant pressed the key corresponding to that square as quickly and accurately as possible. After a key press (irrespective of accuracy) the next cue would immediately appear (response-stimulus interval = 0ms) in a random order. Each practice block included 64 key presses, after which a rest block began and the square outlines turned red again. The task consisted of four practice blocks separated by 15s rest periods. Performance was measured in terms of speed (response time in ms) and accuracy (number of correct key presses).

### Supplementary Results

Supplementary Table 1: Participant Demographics (n=27). Values are mean ± standard deviation. Chronotype data of one participant is missing.

| | |
|---|---|
| Participants Included | 27 |
| Sex (Female) | 13 |
| Age | 23.6 ± 3.5 |
| Edinburgh Handedness | 90.5 ± 15.3 |
| Beck Depression | 1.9 ± 2.2 |
| Beck Anxiety | 2.4 ± 3.0 |
| PSQI | 2.1 ± 1.7 |
| Epworth Sleepiness | 5.9 ± 2.7 |
| Chronotype | 35.7 ± 3.3 |

Supplementary Table 2: MNI coordinates of rsFC ROIs, which were derived from fMRI rsFC literature [2].

| **Seed** | **MNI Coordinates** | | | |
|---|---|---|---|---|
| | **X** | **Y** | **Z** | |
| lANG | -57 | -63 | 17 | DMN |
| rANG | 56 | -63 | 18 | |
| PCC | 5 | -58 | 29 | |
| MPFC | -5 | 35 | -9 | |
| lIPS | -27 | -61 | 50 | DAN |
| rIPS | 26 | -60 | 48 | |
| lFEF | -30 | -9 | 52 | |
| rFEF | 30 | -9 | 55 | |
| rTPJ | 60 | -43 | 16 | VAN |
| rIFG | 42 | 5 | 1 | |
| lTPJ | -54 | -33 | -4 | LANG |
| lIFG | -47 | 14 | 1 | |
| lSMA | -1 | -17 | 55 | MOT |
| lCS | -45 | -17 | 49 | |
| rCS | 45 | -17 | 49 | |
| lS2 | -42 | -13 | 10 | |
| rS2 | 42 | -13 | 10 | |
| lV1V2 | -27 | -81 | -13 | VIS |
| rV1V2 | 27 | -81 | -13 | |
| lMT | -45 | -81 | 4 | |
| rMT | 45 | -81 | 4 | |

Supplementary Table 3: Sleep duration estimated from actigraphy & sleep diaries data for the four days preceding the experimental session (note that due to a lack of compliance to instructions, data of one or more nights are missing for 7 participants; n = 20).

| | **EXP-4** | **EXP-3** | **EXP-2** | **EXP-1** |
|---|---|---|---|---|
| Mean Time Slept (h) | 8.1 ± 1.1 | 8.4 ± 0.8 | 8.3 ± 1.5 | 8.3 ± 0.9 |

Supplementary Table 4: Sleep characteristics from the overnight experimental session EEG recording (data is missing from four participants due to EEG amplifier batteries being depleted of power before the end of the night; n = 23).

| **Experimental Night** | |
|---|---|
| Time in bed (TIB) mins | 475.17 ± 31.99 |
| Total sleep time (TST) mins | 437.06 ± 64.85 |
| Wake after sleep onset (WASO) mins | 25.78 ± 52.61 |
| Sleep onset latency (mins) | 12.32 ± 14.72 |
| REM latency (mins) | 120.69 ± 59.77 |
| Number of awakenings | 11.43 ± 8.03 |
| Efficiency (%) | 92.16 ± 12.78 |
| N1 sleep (mins) | 23.39 ± 11.8 |
| N2 sleep (mins) | 231.76 ± 43.07 |
| N3 sleep (mins) | 75.82 ± 24.56 |
| Total NREM sleep (mins) | 330.97 ± 45.84 |
| REM sleep (mins) | 106.08 ± 32.53 |
| N1 sleep (%TST) | 5.75 ± 4.19 |
| N2 sleep (%TST) | 52.97 ± 5.92 |
| N3 sleep (%TST) | 17.34 ± 5.3 |
| Total NREM sleep (%TST) | 76.07 ± 5.46 |
| REM sleep (%TST) | 23.92 ± 5.46 |

Supplementary Table 5: Values from objective (psychomotor vigilance test – PVT) and subjective (Stanford sleepiness scale – SSS) measure of alertness preceding the training and retest of the MSL task (n = 27).

| | **Training** | **Retest** |
|---|---|---|
| PVT Mean (s) | 0.331 ± 0.045 | 0.326 ± 0.052 |
| PVT Median (s) | 0.311 ± 0.036 | 0.307 ± 0.042 |
| SSS | 3 ± 0.9 | 2 ± 0.8 |

#### 1. Behavior

##### *1.1. Sleep and vigilance*

A repeated-measure ANOVA performed on sleep duration measure extracted from actiwatch and sleep diaries using nights (4) as within-subject factor indicated that sleep duration did not differ across the four nights preceding the experimental session (F(3,57) = 0.546, p = 0.65).

Subjective measures of vigilance (SSS) showed that participants reported to be more alert in the morning retest compared to the nighttime training (SSS training: 2.9 ± 0.86, SSS retest: 1.9 ± 0.77, t(26) = 5.597, p < 0.001). However, such effect was not observed with objective measures of vigilance (PVT training: 0.33s ± 0.04s, PVT retest: 0.30s ± 0.05s, t(28) = 1.38, p = 0.178).

#### 2. Functional Connectivity

##### *2.1. Task-related changes in connectivity*

Exploratory analyses showed task-related decrease in gamma-connectivity within LANG ($F_{(1,20)}$ = 8.17, $p_{uncorr}$ = 0.01) and between LANG-DMN ($F_{(1,20)}$ = 6.264, $p_{uncorr}$ = 0.02) and LANG-VAN ($F_{(1,20)}$ = 7.37, $p_{uncorr}$ = 0.013; see Fig. 3A outside the blue frame in the main text). Additionally, results showed that connectivity in the theta band increased between the pre- to post-task RS sessions within VAN ($F_{(1,20)}$ = 5.06 , $p_{uncorr}$ = 0.036).

##### *2.2. Correlation between connectivity and online gains in performance*

###### *2.2.1. Pre-task (baseline) connectivity*

Negative correlations between baseline delta-band connectivity and online learning were observed between VIS-DAN (r = -0.52, $p_{uncorr}$ = 0.027), VIS-VAN (r = -0.53, $p_{uncorr}$ = 0.025), and VIS-VIS (r = -0.59, $p_{uncorr}$ = 0.009). In the theta band, pre-task connectivity correlated negatively with online learning within the DAN (r= -0.55, $p_{uncorr}$ = 0.017) and VIS (r= -0.64, $p_{uncorr}$ = 0.005), and between VAN-DMN (r= -0.5, $p_{uncorr}$ = 0.034), VIS-DMN (r= -0.59, $p_{uncorr}$ = 0.009), VIS-DAN (r= -0.54, $p_{uncorr}$ = 0.021), and VIS-VAN (r= -0.65, puncorr = 0.004), see main text for motor network correlations. In the beta band connectivity was negatively correlated to online gains within VIS (r = -0.52, $p_{uncorr}$ = 0.028). Gamma band connectivity was found to negatively correlate with online gains within VIS (r = -0.5, $p_{uncorr}$ = 0.035), and between VIS-DMN (r = -0.53, $p_{uncorr}$ = 0.023), VIS-VAN (r = -0.5, $p_{uncorr}$ = 0.036), and VIS-LANG (r = -0.52, $p_{uncorr}$ = 0.027).

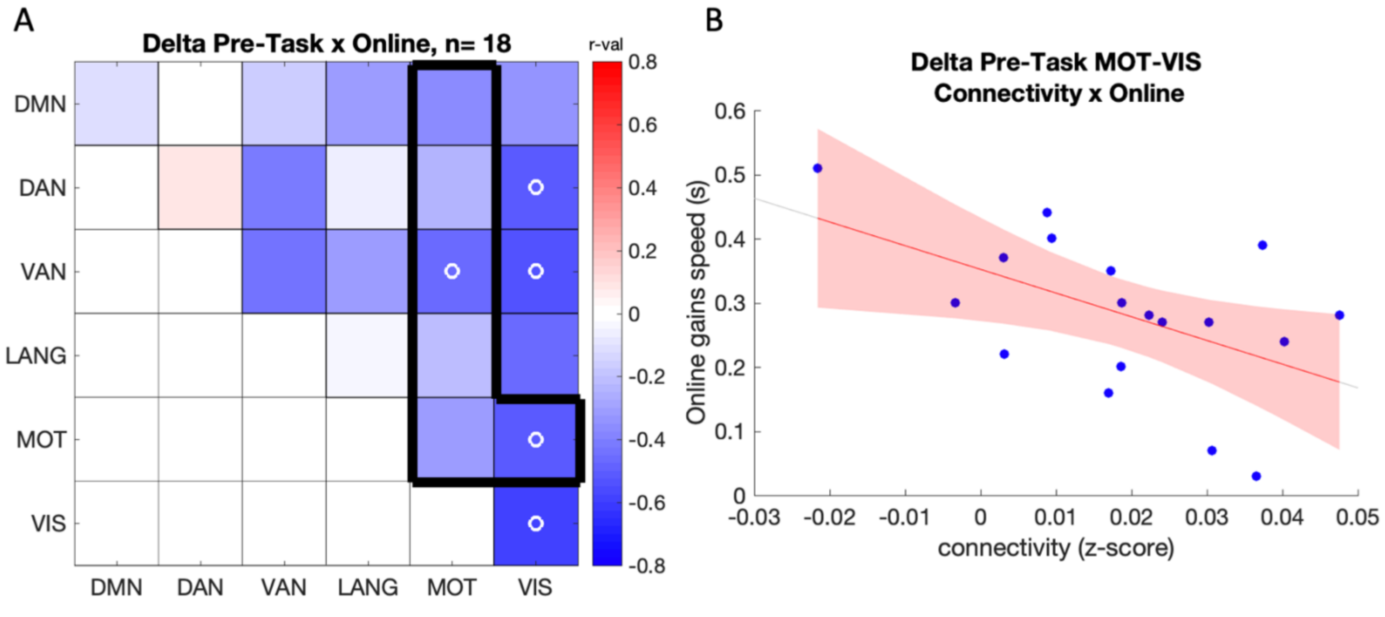


Supplementary Figure 2: A) Correlation between baseline connectivity in the delta band and online gains in performance speed. Color scale depicts r values. Open circles (o) indicate a significant correlation at p < 0.05 uncorrected. None of the above results survived FDR correction for multiple comparisons. B) Scatter plot displaying the correlation between online gains in speed and baseline connectivity in the delta band between the motor and visual networks (n = 18).

*2.2.2. Post-task connectivity*

There were no significant correlations between post-task rsFC and online gains in performance in the motor network pairs and frequency bands of primary interest. However, exploratory analyses revealed that higher theta post-task connectivity between MOT-DMN (r = -0.56, $p_{uncorr}$ = 0.016), MOT-VAN (r = -0.51, $p_{uncorr}$ = 0.032) and MOT-VIS (r = -0.57, $p_{uncorr}$ = 0.014) was negatively correlated to online gains in performance. Similar negative correlations were observed in the delta band in network pairs of no interest (i.e, within the DMN (r = -0.7, $p_{uncorr}$ = 0.001), VAN (r = -0.54, $p_{uncorr}$ = 0.022), and VIS (r = -0.52, $p_{uncorr}$ = 0.028), and between the DMN-VAN (r = -0.68, $p_{uncorr}$ = 0.002), DMN-VIS (r = -0.68, $p_{uncorr}$ = 0.002), LANG-VAN (r = -0.58, $p_{uncorr}$ = 0.011), and LANG-VIS (r = -0.56, $p_{uncorr}$ = 0.016); supplemental Fig. 3).

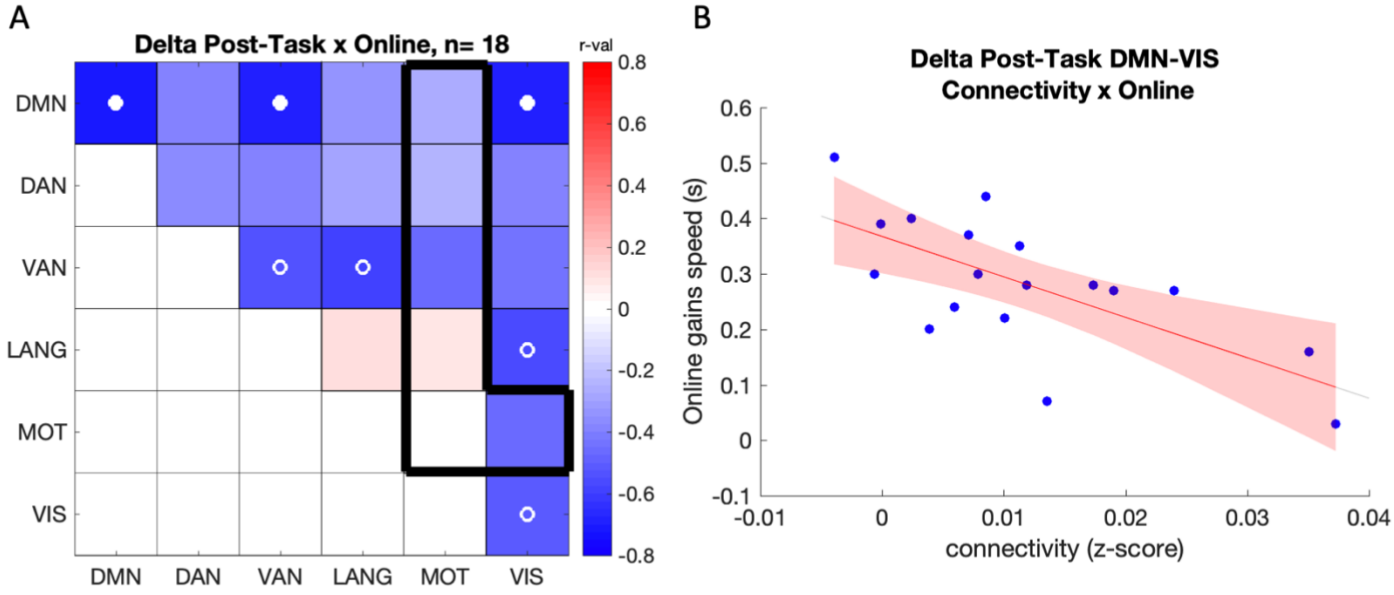


Supplementary Figure 3: A) Correlation between post-task connectivity in the delta band and online gains in speed. Color scale depicts r values. Open circles (o) indicate a significant correlation at $p < 0.05$ uncorrected, while (•) marks $p < 0.05$ FDR corrected across all 21 comparisons. B) Scatter plot displaying the correlation between online gains in speed and baseline connectivity in the delta band between the motor and visual networks (n = 18).

*2.2.3. Task-related changes in connectivity*

Exploratory analyses revealed that task-related decreases in gamma band connectivity between VIS-DMN (r = 0.49, $p_{uncorr}$ = 0.04), VIS-VAN (r = 0.49 , $p_{uncorr}$ = 0.041) and within VIS (r = 0.48 , $p_{uncorr}$ = 0.047) correlated positively with online gains in performance (Supplementary figure 4). This relationship was also observed in the theta band between VIS-VAN (r = 0.51, $p_{uncorr}$ = 0.03).

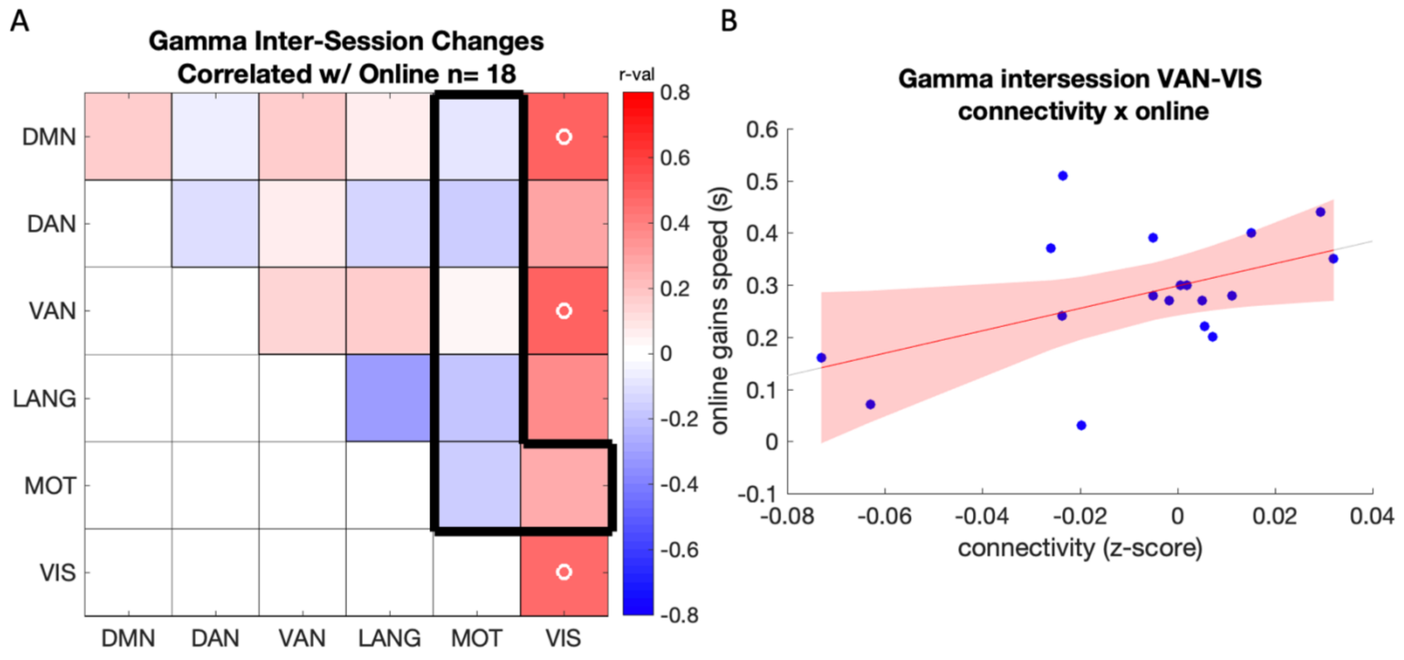


Supplementary Figure 4: A) Correlation between online gains in performance speed and inter-session (post – pre task) connectivity changes in the gamma band (n = 18). Color scale depicts r values. Open circles (o) indicate a significant correlation at $p < 0.05$ uncorrected. None of the above results survived FDR correction for multiple comparisons. B) Scatter plot displaying the correlation between online gains in speed and intersession changes in connectivity in the gamma band between the default mode and visual networks (n = 18).

##### *2.3. Correlation between connectivity and offline gains in performance*

###### *2.3.1. Pre-task (baseline) connectivity*

A positive relationship between pre-task connectivity and offline gains in speed was observed in the theta band within VAN ($r = 0.52$, $p_{uncorr} = 0.049$) and within LAN network ($r = 0.61$, $p_{uncorr} = 0.016$). Additionally, in the alpha band a negative relationship was found between overnight improvements in speed and connectivity between VAN-VIS ($r = -0.534$, $p_{corr} = 0.04$). Beta-band connectivity negatively correlated to offline gains between VIS-DMN ($r = -0.56$, $p_{uncorr} = 0.03$), VIS-DAN ($r = -0.62$, $p_{uncorr} = 0.014$), VIS-VAN ($r = -0.51$, $p_{uncorr} = 0.05$), and VIS-LANG ($r = -0.66$, $p_{uncorr} = 0.008$).

###### *2.3.2. Post-task connectivity*

In the beta band a negative relationship was found between overnight changes in performance and post-task connectivity between LANG-VIS ($r = -0.547$, $p_{uncorr} = 0.035$) and LANG-VAN ($r = -0.516$, $p_{uncorr} = 0.049$).

###### *2.3.3. Task-related changes in connectivity*

A negative relationship was found between offline gains in performance and task-related connectivity increases within the language network in the theta band ($r = -0.68$, $p_{uncorr} = 0.005$). In addition to the results reported in the main text, task-related changes to beta-band connectivity positively correlated with offline gains between VIS-DAN ($r = 0.67$, $p_{uncorr} = 0.006$), and VIS-VAN ($r = 0.54$, $p_{uncorr} = 0.036$).
